# Imitation Predicts Social Favor in Adolescent Rhesus Macaques (*Macaca mulatta*)

**DOI:** 10.1101/208777

**Authors:** Jordan Anderson, Erin L Kinnally

**Affiliations:** Duke University, Department of Evolutionary Anthropology; University of California Davis, Department of Psychology and Animal Behavior Graduate Group; California National Primate Research Center, Davis, CA

**Keywords:** Imitation, Macaca mulatta, Social Relationships, Juveniles

## Abstract

**Objectives:** Imitation is a highly conserved component of animal behavior with multifaceted connections to sociality across taxa. One intriguing consequence of imitation in primates is that it promotes positive social feedback from the imitated toward the imitator. This suggests that imitation in primates may facilitate positive social interactions, but few studies have tracked imitation in socially housed primates. Here, we designed a novel ethogram to characterize imitation between conspecifics, to better understand whether imitation is associated with affiliation between primates in a semi-natural setting.

**Materials and Methods:** In this study, 15 juvenile rhesus macaques (*Macaca mulatta*) were observed at the California National Primate Research Center. Using focal sampling, frequencies of imitative events (e.g. following, postural mimicry, etc.) by the focal were observed over a course of 12 weeks. In separate observations, focal social behavior (e.g. aggression, play, etc.) was also observed.

**Results:** Subjects that exhibited higher degrees of imitation were not necessarily more prosocial, but, consistent with our hypothesis, they received significantly more play overtures from social partners (p<.01). In addition, imitation rates generally decrease with age.

**Conclusions:** Together, these results suggest that imitation is associated with receiving positive social behavior in a complex, semi-natural setting in primates, and that imitation may be more common in adolescence as opposed to adulthood. These preliminary results in a small sample set represent an important step in characterizing imitation in context of social interactions during development. Tracking these behaviors over time will elucidate whether imitation is directly recruiting these positive social interactions, as has been demonstrated in captivity.

## INTRODUCTION

The social environment of primates is complex, and the ability to navigate it to develop strong social relationships has adaptive consequences. Primates with strong social connectivity live longer, and social connectedness may increase an individual’s access to mates, lifespan, or offspring survival, thereby increasing its potential reproductive output (Silk et al., 2003; Silk, 2007; Silk et al., 2010; Archie et al., 2014). Yet there are certain facets of primate social behavior which may have an underexplored impact on these types of social success. In particular, imitation as a behavioral construct has been of interest to researchers for decades.

Imitation may take many forms, and as such has taken on a range of definitions and characterizations. For example, researchers have been interested in the timing of imitative events (e.g. synchronous versus delayed imitation), the nature of the imitation (e.g. cognitive versus motor), as well its differentiation from other mirroring behavior (e.g. emulation and copying), among other things (see brief overview by Zentall, 2003). However broadly speaking, imitation may simply be considered the matching of one’s own behavior with that of another individual, regardless of the function or mechanism (Chartrand and Bargh, 1999; Ferrari et al., 2006). From this broader perspective, there is strong evidence that numerous primates can and do imitate in a variety of social contexts and in a variety of ways (Voelkl & Huber, 2000; Ross et al., 2008; Byrne 2006, Subiaul et al., 2004; Whiten et al., 1996).

In great apes, imitation has often been studied in the context of social learning, where the ability to match one’s behavior to a conspecific may be an important means of skill acquisition (Nagell et al., 1993; Whiten et al., 1996; Miller & Dollar 1941). In recent years, a greater focus has been placed on understanding whether imitation plays a role in pro-social interaction.

Humans, for example, non-consciously imitate the behavior (e.g. mannerisms and posture) of social partners (Chartrand & Bargh, 1999). This has positive social consequences for the imitator: imitated individuals reported higher regard (measured through later self-reported ratings) towards their imitator versus a confederate non-imitative transaction partner (Chartrand & Bargh, 1999). Intriguingly, similar phenomena have been reported between distantly related primate species, perhaps highlighting its adaptive significance. In captivity, imitation of both conspecifics (Ross et al., 2008) and human observers (Byrne, 2006), has been observed in great apes, although it is largely unknown whether these forms of imitation have favorable long-term social consequences. Adolescent and young adult brown capuchins (*Cebus apella*) spent more time gazing at, maintaining proximity with, and engaging in a token exchange task with imitators over non-imitators, suggesting a preference for imitators (Paukner et al., 2009). In geladas (*Theropithecus gelada*), rapid facial mimicry during play-bouts positively predicts the length of the play interaction (Mancini et al., 2013). Together these studies strongly suggest that imitation may impact numerous aspects of one’s social environment. However, in general, the social consequences of imitation in semi-natural settings remains poorly understood.

Imitation may be trait-like, in that some monkeys imitate significantly more than others, but it may also shift across development to serve evolving social strategies. In newborn infant rhesus macaques (*Macaca mulatta*), not all monkeys imitate human experimenters (Simpson et al., 2016). This trait may contribute to later social competence: macaque imitators that imitate human experimenters exhibit more pro-sociality across development, including more gaze following of a human experimenter, more eye tracking of a conspecific’s face, and discrimination of familiar *vs*. novel human experimenters, compared with non-imitators (Paukner et al., 2014; Simpson et al., 2013; Simpson et al., 2016). The role of imitation later in life is similarly not well known in monkeys. In one study of adult capuchins (*Cebus apella*) under experimental conditions, subjects given foraging options tended to choose the same option as a familiar conspecific, showing that imitation may continue to play a role in adult capuchin life (Bonnie & De Waal, 2007).

Thus, primates appear to imitate in a variety of contexts. However, it remains unknown whether social imitation is associated with pro-sociality in naturally-occurring social life. This is due, in part, to the difficulty of characterizing imitation in natural/semi-natural settings. Given the demonstrated importance of developing and maintaining strong social bonds in primates, exploring the relationship between imitation and affiliation remains critical to understanding the evolution of these processes and their role in primate social life.

To help bridge these gaps, this study aimed to test whether there is a relationship between imitation and affiliation in semi-naturalistically housed rhesus macaques (*Macaca mulatta*). If imitation facilitates the development and maintenance of social bonds, we would expect individuals that exhibit higher degrees of imitation to have more affiliative social interactions. To test this, we observed macaques in a semi-natural, captive social setting over a 12-week time course. We then characterized the prevalence of imitation in socially housed primates, and next, we addressed whether imitation is associated with greater social affiliation. By observing individuals ranging from adolescence (2 years old) to early adulthood (4 years old) we also aimed to investigate how propensity to imitate may differ between these developmental stages.

## METHODS

### Subjects

Subjects (n=15) were two to four-year-old (mean=2.97) juvenile rhesus macaques (*Macaca mulatta*; 8 males, 7 females) living in one of three semi-naturalistically housed social groups at the California National Primate Research Center (CNPRC) in Davis, California. Seminatural housing covers half an acre, and is enclosed by chain link fencing. Social groups included the range of age/sex groups, matrilineal social hierarchy structure and at least six distinct matrilines. Social rank is assessed on a monthly basis by trained CNPRC staff as part of normal colony procedure. Data were collected in all field corrals for at least 30 minutes on a bi-weekly basis, totaling at least one hour of data collection per month. Data were collected with scan sampling to record dyadic displacement interactions between individuals. This data is entered into a hierarchy grid to analyze for the best hierarchical configuration in each cage. Separate rank hierarchies are determined for males and females. We calculated social rank as the absolute rank/number of animals of the same sex in the social group. Ranks ranged from the 25% percentile (more highly ranked animals) to the 97th percentile (lower ranked animals; mean 58%). Animals had access to water *ad libitum* and are fed monkey chow biscuits (Purina, Inc) twice daily, once in the morning and once in the afternoon.

### Social behavior observations

Focal observations were conducted once per week for five minutes between the hours of 0800 and 1200 for 12 weeks (50 minutes of total data were collected for each individual over ten observations) by the same male observer. Observations included a combination of 15-second scans for social state. Social states included proximity (within 1 meter) from another individual and physical contact with another individual. All occurrences of social transactions were also observed. Social transactions were adapted from a transactional coding scheme described previously (Kinnally, 2014). A new transaction was recorded when a pair of animals changed their state of interaction. Social transactions were recorded when the focal either received or engaged in affiliation (proactive contact), aggression (agonistic contact), or play (patterns of behavior resembling those used in serious functional contexts, like biting, fighting, fleeing … but used in activities that appear to have no obvious immediate benefits to the players; adapted from Worch, 2012). The response was then recorded, as well as the focal’s role as either the initiator or recipient of the transaction. Possible responses to an overture included play, affiliation, neutral, resistance, and aggression (see Table 1, Section 1 for definitions; see Figure 1 for frequencies observed per observation).

**Figure 1.**
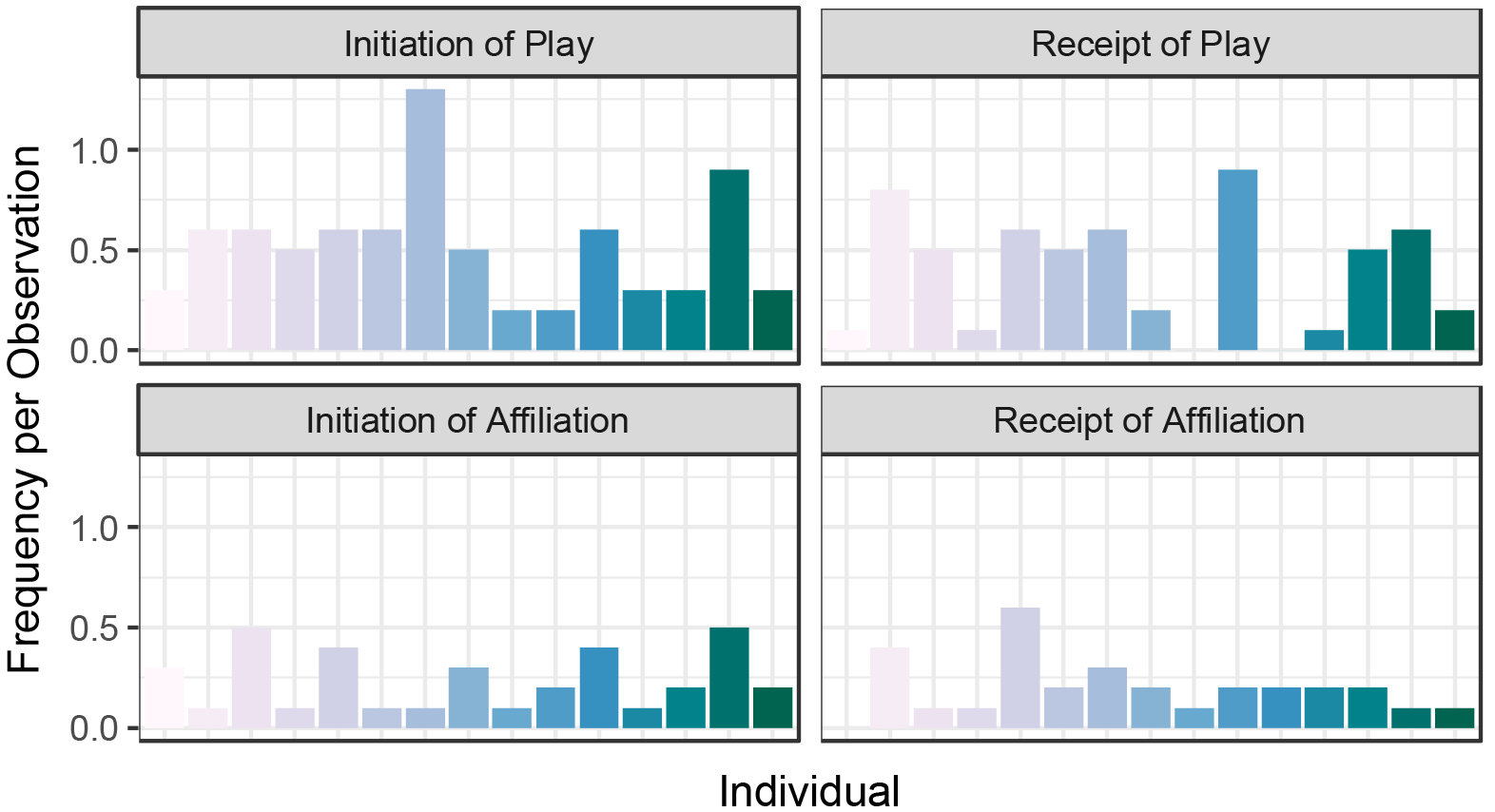
The frequency per observation of the four most common social behaviors, the initiation of play, the receipt of a play overture, the initiation of affiliation, and the receipt of an affiliative overture for each individual. Colors represent different individuals.

**Table 1.** Ethogram of Social and Imitation Behaviors.

| **Section 1: Social Behaviors (adapted from Kinnally et al., 2014)** |  |
| --- | --- |
| **Play** | Patterns of behavior resembling those used in serious functional contexts—fighting, fleeing, traveling, and mounting—but used in activities that appear to have no obvious immediate benefits to the players (adapted from Worch, 2012). |
| **Affiliation** | Any form of prosocial (non-aggressive) physical contact between two individuals (excluding play behavior). |
| **Neutral** | No response, positive or negative, to an overture (e.g. no change in behavior). |
| **Resistance** | The active denial of an overture by leaving the area of the initiator. |
| **Aggression** | Threatening, biting, chasing, scratching, flattening (pressing into the ground), dragging, pushing away, or grabbing. |
| **Section 2: Imitation Behaviors** |  |
| **Following** | The direct following behind another individual. This was distinguished from chasing and play behavior in that these behaviors resulted in agonistic and prosocial interactions respectively following a period of tracking. In contrast, for a transaction to be considered following, tracking ceased for >3 seconds prior to any other behavioral transactions occurring. |
| **Object Exploration** | Interaction with the same non-novel object (e.g. Cage mirrors, toys, etc.). |
| **Foraging** | The consumption of the same non-novel food item by only two individuals outside structured feeding schedules. |
| **Auto-grooming** | Any form of self-grooming by an individual shortly after (<5 seconds) a nearby conspecific. |
| **Postural** | The repositioning of one's posture to mirror the atypical posture of a nearby static individual. The most common postures: quadrupedal standing and sitting were excluded from this form of imitation. Examples of atypical posture include, but are not limited to, resting feet on fence at chest-level while sitting, laying fully supine or fully prone, or bipedal standing. |
| **Grooming** | The grooming of the same third party individual by two individuals either simultaneously or in short succession (<5 seconds). |
| **Yawning** | Yawning by an individual shortly after that of a nearby conspecific. |

### Imitation observations

Subjects were observed for imitative behavior over the same 12-week time period as social behavior observations were conducted, but on different days. Imitative data were collected during 5-minute focal observations conducted twice per week, for a total of 14 observations per subject. Imitation data were collected on separate days to prevent dependence of social and imitation observations. Imitation was considered to be engaging in behaviors mirroring that of another individual in the social group within a pre-specified temporal, spatial, and social context. The criteria for imitation were fourfold. First, an imitator must be within three meters of the imitated individual and within sight of each other. Second, we included only events in which the pair were the only individuals within three meters engaging in the target behavior. We included this criterion to preclude the possibility that an animal was “joining a crowd” engaging in some collective behavior. Thirdly, the imitation had to occur no later than five seconds after the end of the imitated behavior. Finally, imitation behaviors fell into one of the following categories: exploring the same objects (Object Exploration), foraging in the same area as a conspecific outside typical foraging time (Foraging), moving in the same direction (Following), postural changes that mimicked that of a nearby conspecific (Postural), self-directed grooming (Autogrooming), third party grooming (Grooming), and sequential yawning (Yawning; see Table 1, Section 2 for definitions; see Figure 2 for frequencies observed).

**Figure 2.**
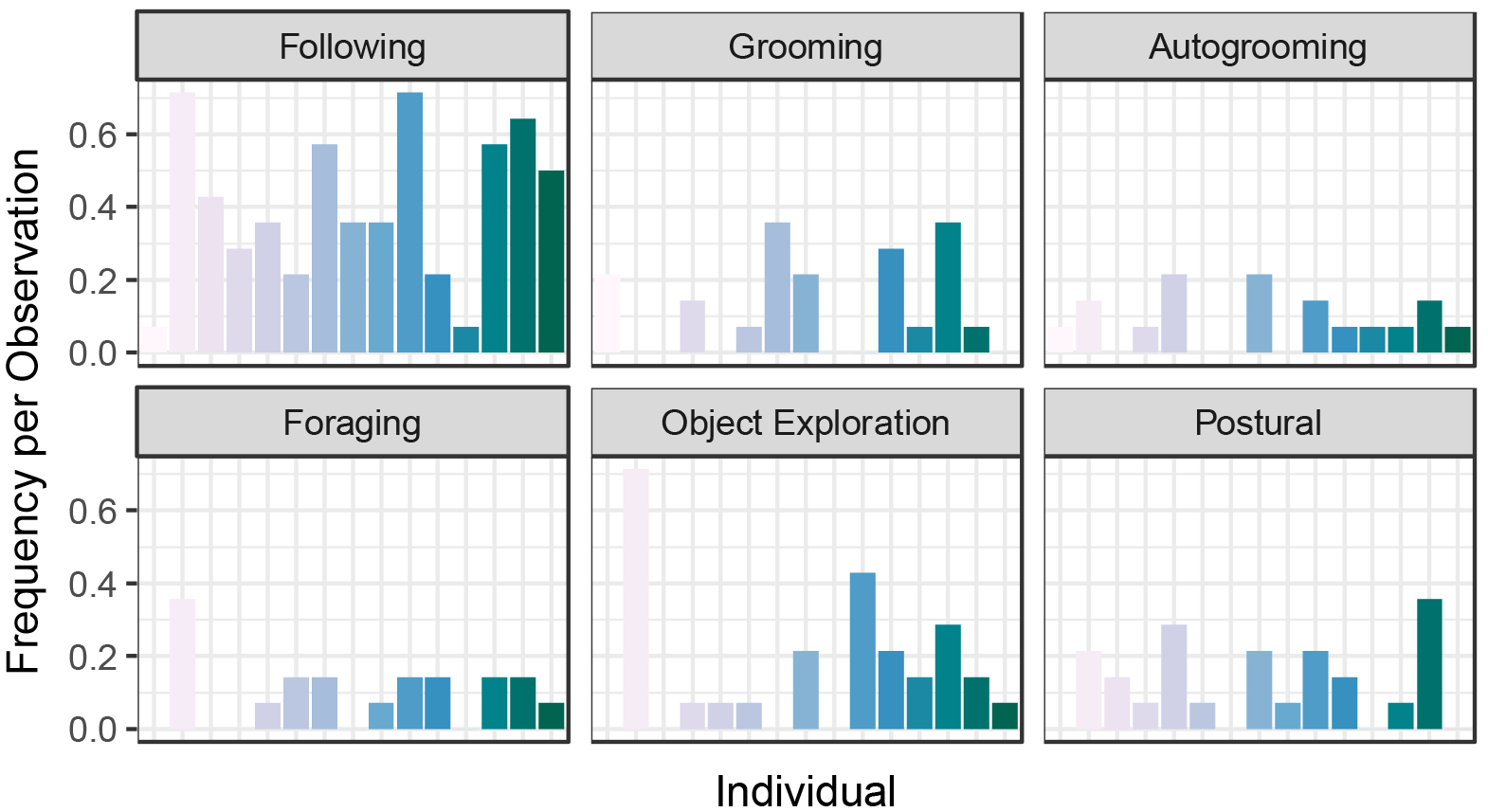
The frequency per observation of the six putative imitative behaviors for each individual. Colors represent different individuals.

Although imitation has not been directly characterized in social contexts, previous literature on imitation in captive settings provided expectations for ways in which imitation may be manifested in a semi-natural environment. For example, due to the fact that foraging decisions are subject to copying by conspecific brown capuchins (Bonnie & De Waal, 2007), we might predict that some juveniles are likely to be seen mimicking the movement of others, or to be frequently found contemporaneously foraging with others outside their normal eating schedule (Following; Foraging). Rhesus macaques have also demonstrated the ability to mimic how conspecifics interact with an object (Subiaul et al., 2004). As such, juveniles may be expected to show similar patterns of mimicry with objects in their captive environment (e.g. enrichment; Object Exploration). Even more, humans have been shown to imitate mannerisms of interaction partners (Chartrand and Bargh, 1999). From this, we might predict that juveniles often re-orient there position to match that of nearby conspecific, or to engage in auto-grooming if they observe another individual engaging in this behavior (Postural; Autogrooming). This should not be considered to be an exhaustive list of possible imitative events in juvenile rhesus macaques. Rather these were chosen as behaviors that may be logical extensions of previously demonstrated imitation behaviors that could be feasibly collected given our constraints and were frequently observed during ethogram development.

### Data analysis

All data analysis was done with the R statistical software (v 3.3.0) using the R Studio integrated development environment (R Core Team, 2016; R Studio Team 2015). First, we determined the relations among imitative behaviors using Cronbach’s Alpha, an indicator of internal reliability. Behaviors with high inter-reliability (Cronbach’s Alpha >.70) were entered into a factor analysis (principal components analysis, promax rotation, “prcomp” function in the “stats” package) to determine whether latent dimensions of imitative behavior could be determined. We next examined whether demographic factors, such as age, sex, or social rank, predicted these dimensions using analysis of variance. Following this, we were interested in detecting whether aspects of social behavior were predicted by imitation dimensions. We focused on the most common transaction types: play and affiliation. Other transaction types, such as aggression and neutral interactions, were relatively rare (median < 0.2 events per observation). We examined the relationship between imitation and rates of play initiated and received, as well as affiliation initiated and received, using multiple regression. For each model, demographic factors were included in the model and removed if non-significant using ANOVA tests (“aov” function in the “stats” package). Significance was set at p <.05.

## RESULTS

Cronbach’s alpha analysis revealed a high degree of reliability (Cronbach ‘s alpha = .775) among five of the imitation behaviors (Object Exploration, Following, Foraging, Autogrooming, and Postural imitation; see Figure 2 for rates of observed imitation). Grooming a third party did not correlate with these behaviors and was therefore removed from subsequent analyses. Factor analysis and principal components analysis (PCA) of the five imitation behaviors revealed two underlying factors in imitation behavior. The first factor included following, foraging, and object exploring, and explained 43.5% of the variation in imitation. We refer to this factor as “Environment-directed Imitation”, because it includes imitative behaviors that include interacting with the environment. The second factor included postural and autogrooming imitative events, and explained 35.7% of the variance. We refer to this factor as “Self-directed imitation” because these behaviors are self-directed, rather than outwardly directed. Enviroment-directed and self-directed imitation scores were not significantly correlated (p=.156, adjusted r^2^=.083; see Figure 3 for the relationship between these factors and each imitative behavior). The most frequent imitative behaviors observed were following and object exploration and the rarest was autogrooming (Figure 2)

**Figure 3.**
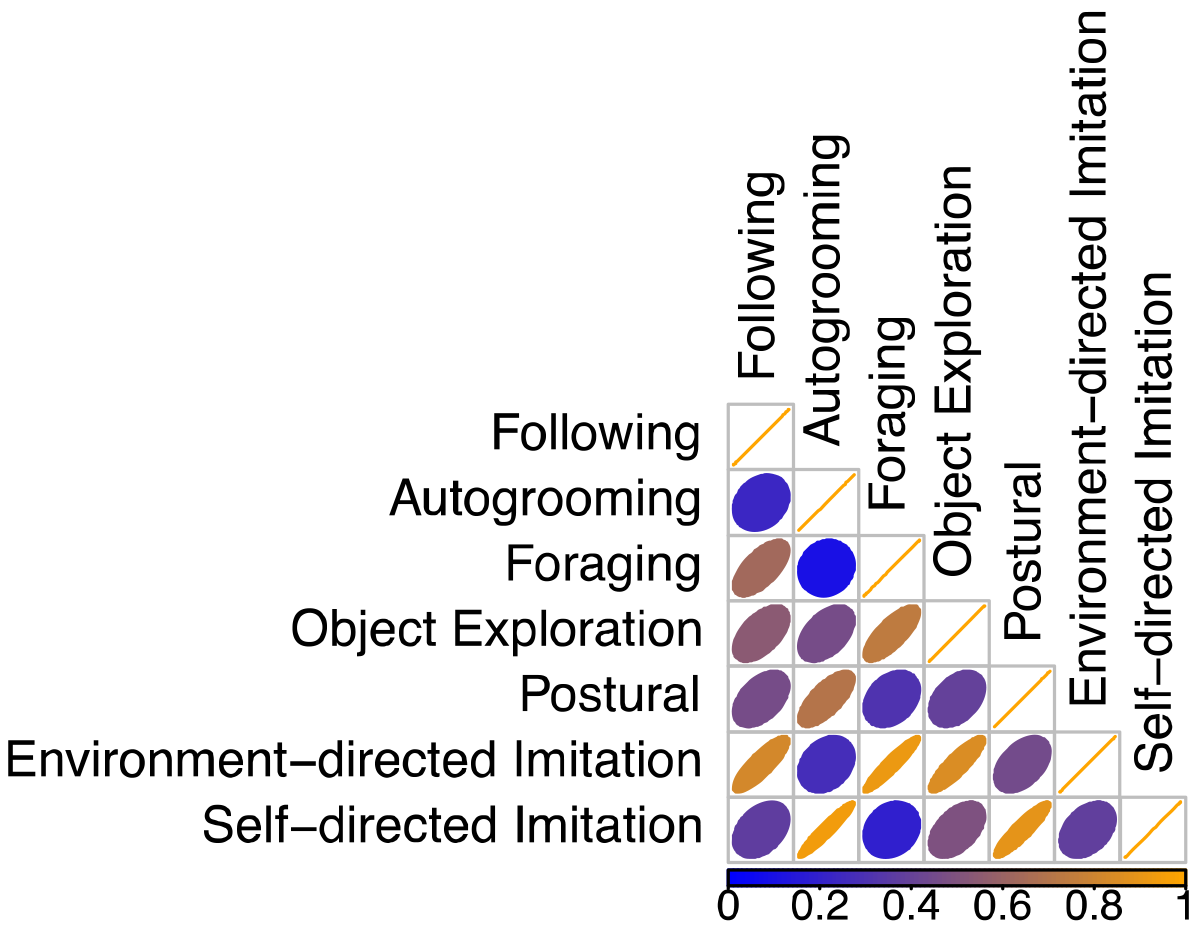
The first two factors from the principle component analysis versus the five retained imitative behaviors- Following, Foraging, Object Exploration, Auto-grooming, and Postural imitation.

Testing for the effects of demographic factors on imitation, we found no significant effect of social group, sex, or social rank on either Enviroment-directed or self-directed imitation (all p >.05). We did, however find that age significantly affected rates of Environment-directed, but not self-directed, imitation (p=.009 and p>.05 respectively). The effect was such that three year olds performed the most social imitation, and four year olds the least. Similarly, cage, sex, and social rank explained no significant variation in any of the social behaviors (all p>.05). As expected, age significantly correlated with the initiation of play bouts, and was correlated at trend level with the receipt of play (p=.035 and p=.085 respectively).

Finally, imitation factor scores predicted aspects of social behavior, even when statistically controlling for the effects of age. Environment-directed imitation scores predicted the rate at which subjects received play overtures from conspecifics (p=.002, adjusted r^2^=.52; Figure 4), but not the rate at which they received affiliation (p>.05). Environment-directed imitation scores did not predict the rate at which focals initiated play (p>.05; Figure 5) or affiliation (p>.05). Additionally, self-directed imitation did not significantly predict any aspect of social behavior (p> .05), although it did predict at trend level the frequency of receiving an affiliative overture (p=.085).

**Figure 4.**
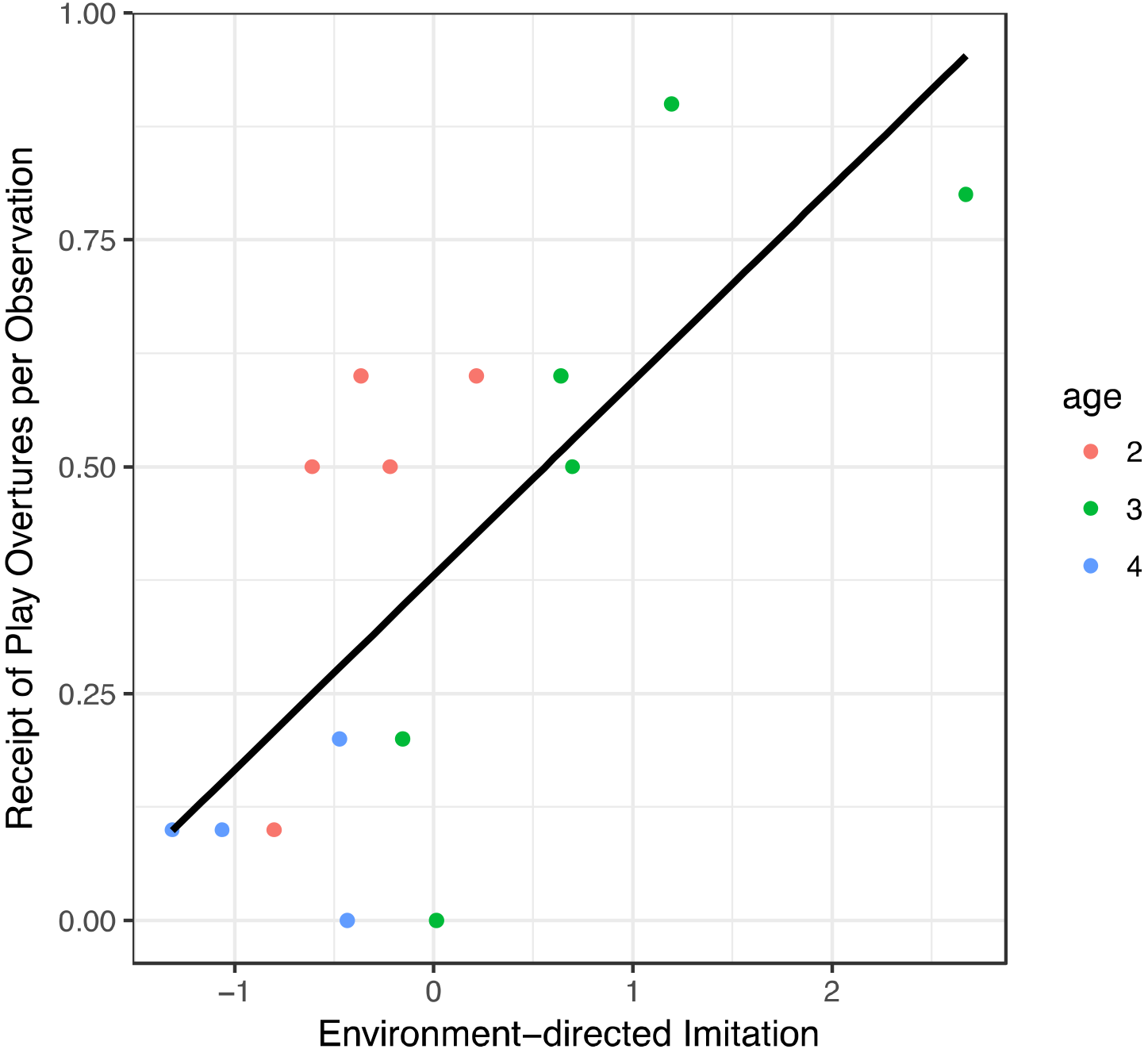
The relationship between environment-directed imitation factors and the frequency of the receipt of play overtures per observation, r^2^=.5247: p=0.002.

**Figure 5.**
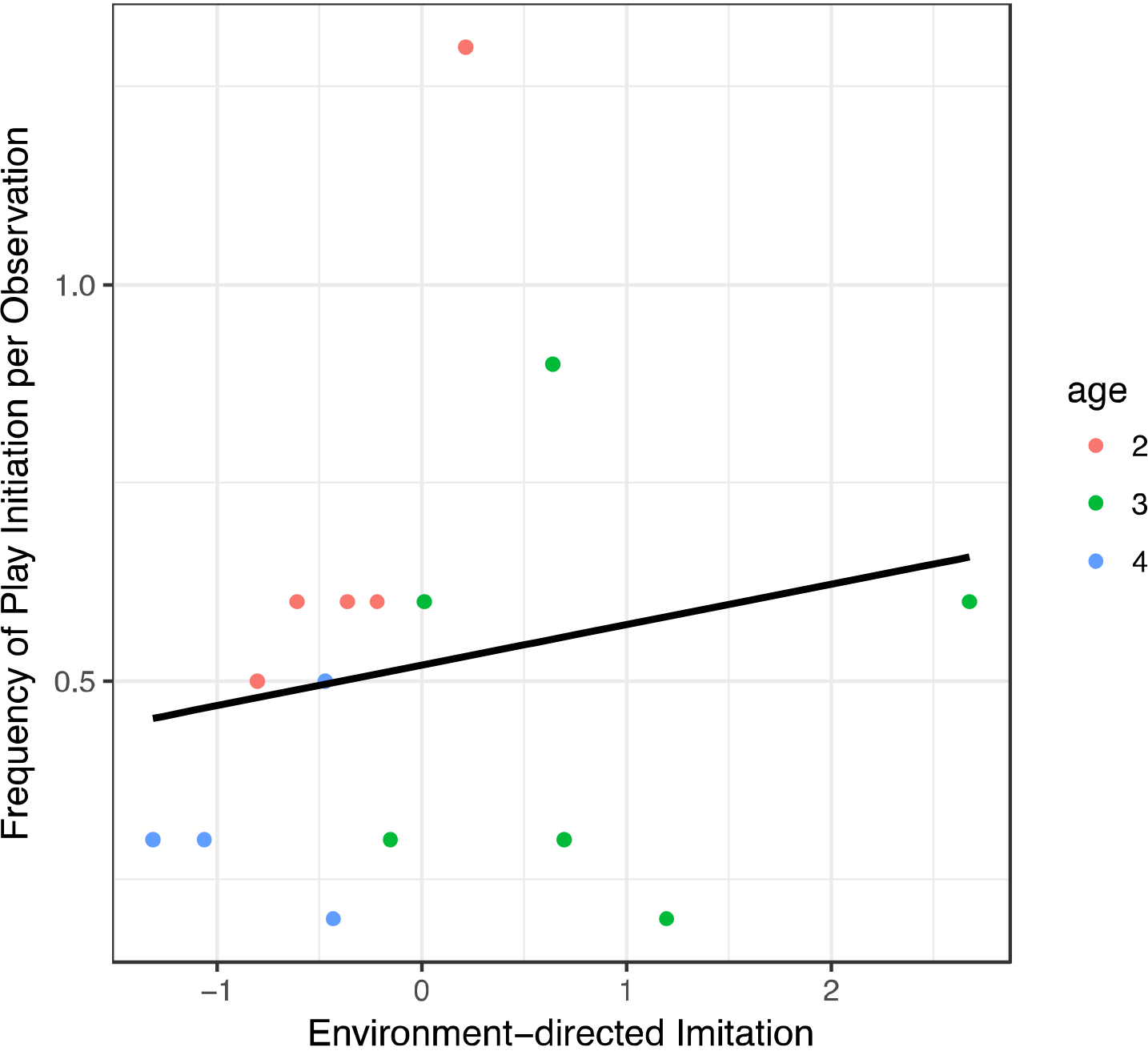
The relationship between environment-directed imitation factors and the frequency of initiating play per observation, p>.2.

## DISCUSSION

Concordant with previous studies, we find that rates of imitation positively predicted one aspect of affiliation in semi-naturalistically housed primates. Individuals exhibiting greater rates of imitation received significantly more play overtures from conspecifics. However, the degree of imitation exhibited did not predict propensity to initiate play or affiliation. These data suggest that imitative capacity or motivation varies between individual macaques. It also suggests that imitation is not necessarily linked with social motivation, but may predict positive social favor.

To our knowledge this is the first study to observe naturalistic types of imitation in rhesus macaque juveniles. We detected two general dimensions of imitative behavior, which we termed environment-directed and self-directed imitation. Environment-directed imitation consisted of mimicking another animal by following them or imitating their foraging and object exploration behaviors. Self-directed imitation included mimicking auto-grooming and posture. It is notable however that these two types of imitation are not significantly correlated. Demographic factors like social rank and sex did not predict imitation rates in the present study. This too suggests that imitative propensity may be a unique individual trait.

Play was the most frequent type of social transaction observed for juveniles. Because imitators received more play overtures, this means that environment-directed, but not self-directed, imitation rates predicted one of the most frequent, and possibly most important aspect of social interaction at this developmental stage. A limitation of this study is that we cannot know whether imitation is linked with a third aspect of social behavior that in fact enhances social favor, or is subject to reverse causality. If there were a second correlated trait with imitation that predicted social favor, we would expect individuals that both environmentally and self-directedly imitate might share this trait and receive some degree of social favor. However we find no evidence for this here, as self-directed imitation did not significantly predict the receipt of play overtures.

Like many behaviors, the developmental timing of imitation in the social realm may be important. We found here that 4 year olds engaged in significantly less imitation than 2 and 3 year olds. However, due to the fact that our observations are cross-sectional, rather than longitudinal, we cannot say how individual’s propensity to imitate may change over their lifespan. However, strong social bond acquisition early in life has proven longitudinal consequences (Weinstein & Capitanio, 2008; Weinstein & Capitanio, 2012), thus the demonstrated prevalence of imitation at critical early life periods may be linked to positive consequences later in life.

## LIMITATIONS

Although this study aimed to bridge numerous gaps in our knowledge of the social implications of imitation, understanding the limitations of this study design may further aid future studies in addressing gaps left here. Obviously, our sample size was small. Future studies should examine the relations between imitation and sociality in a larger cohort. Most importantly, imitation of this nature is the reflection of a dyadic relationship, but we considered only rates of imitation of each subjects and rates of social interactions. We did not track whether imitators imitated some individuals more than others (e.g. individuals that have shown them social favor previously). Thus, we cannot be sure of the direction of the relationship between imitation and social favor. It is possible that animals imitate animals with whom they have preexisting affiliative bonds. Studies focusing on dyadic relationships will better demonstrate the causal effect of imitation on affiliation, and how imitation varies across relationships within an individual. Even more, the relationship between imitation and affiliation is likely one that develops over time. Thus cross-sectional studies such as these are insufficient to show any true causal relationship. Tracking both social interactions and imitative behavior over time will allow us to directly test for a causal relationship. The behaviors presented here as imitation are an initial attempt at characterizing imitation in semi-natural settings. Future research expanding the imitation ethogram will serve to give us a fuller picture of the forms imitation may take. Finally, the confinement of semi-natural conditions may promote higher levels of interactions between individuals, so to understand the true magnitude of this effect, an extension to wild-ranging systems remains crucial.

## ACKNOWLEDGEMENTS

This work was funded by the California National Primate Research Center Base Grant (P51 OD0011107) and a UC Davis Provost’s Undergraduate Research Fellowship (to JA). We thank anonymous reviewers for their helpful comments during previous drafts of this manuscript.

